# SARS-CoV-2 infection enhancement by Amphotericin B: Implications for disease management

**DOI:** 10.1101/2024.11.07.622419

**Authors:** Dung Nguyen, Stephen M. Laidlaw, Xiaofeng Dong, Matthew Wand, Amanda Horton, Mark Sutton, Julia Tree, Rachel Milligan, Maximilian Erdmann, David Matthews, Andrew D. Davidson, Khondaker Miraz Rahman, Julian A. Hiscox, Miles Carroll

## Abstract

Severe coronavirus disease 2019 (COVID-19) patients who require hospitalisation are at high risk of invasive pulmonary mucormycosis. Amphotericin B (AmB), which is the first line therapy for invasive pulmonary mucormycosis, has been shown to promote or inhibit replication of a spectrum of viruses. In this study, we first predicted that AmB and Nystatin had strong interactions with severe acute respiratory syndrome coronavirus 2 (SARS-CoV-2) proteins using in silico screening, indicative of drugs with potential therapeutic activity against this virus. Subsequently, we investigated the impact of AmB, Nystatin, Natamycin, Fluconazole and Caspofungin on SARS-CoV-2 infection and replication *in vitro*. Results showed that AmB and Nystatin actually increased SARS-CoV-2 replication in Vero E6, Calu-3 and Huh7 cells. At optimal concentrations, AmB and Nystatin increase SARS-CoV-2 replication by up to 100- and 10-fold in Vero E6 and Calu-3 cells, respectively. The other antifungals tested had no impact on SARS-CoV-2 infection *in vitro*. Drug kinetic studies indicate that AmB enhances SARS-CoV-2 infection by promoting viral entry into cells. Additionally, knockdown of genes encoding for interferon-induced transmembrane (IFITM) proteins 1, 2, and 3 suggests AmB enhances SARS-CoV-2 cell entry by overcoming the antiviral effect of the IFITM3 protein. This study further elucidates the role of IFITM3 in viral entry and highlights the potential dangers of treating COVID-19 patients, with invasive pulmonary mucormycosis, using AmB.

**Importance:** AmB and Nystatin are common treatments for fungal infections but were predicted to strongly interact with SARS-CoV-2 proteins, indicating their potential modulation or inhibition against the virus. However, our tests revealed that these antifungals, in fact, enhance SARS-CoV-2 infection by facilitating viral entry into cells. The magnitude of enhancement could be up to 10- or 100-fold depending on cell lines used. These findings indicate that AmB and Nystatin have the potential to enhance disease when given to patients infected with SARS-CoV-2 and therefore should not be used for treatment of fungal infections in active COVID-19 cases.

## Introduction

SARS-CoV-2 is a member of the family *Coronaviridae*, genus *Betacoronavirus*, species *Severe acute respiratory syndrome*-*related virus*. The virus enters cells via endocytosis or via plasma membrane fusion following binding to its receptor, angiotensin-converting enzyme 2 (ACE2) [1–4]. It has been reported that the preference of entry pathway used by SARS-CoV-2 is determined by transmembrane serine protease 2 (TMPRSS2) expression on target cells [5, 6]. In cells expressing TMPRSS2 (for example Calu-3), SARS-CoV-2 uses a fast route (plasma membrane fusion) to enter cells. In cells with insufficient TMPRSS2 expression (Vero E6, Huh7.5), SARS-CoV-2 relies on a slow pathway (clathrin-mediated endocytosis via cathepsin L) for infection [7]. While TMPRSS2 activity at the cell surface is pH independent, cathepsin L requires a low pH environment to be activated [8]. Although the plasma membrane fusion route is more efficient and results in more viral particle production for some coronaviruses [9, 10], the recent SARS-CoV-2 Omicron variant has evolved to be less dependent on TMPRSS2 for cell entry [11].

As of the end of October 2024, SARS-CoV-2 has caused more than 700 million COVID-19 cases and 7 million deaths worldwide (https://www.worldometers.info/coronavirus/). About 5% of COVID-19 patients experience severe infection and require intensive care unit admission [12, 13]. These patients, particularly those with diabetes or receiving corticosteroids as treatment for COVID-19, are at high risk of secondary infections including invasive pulmonary mucormycosis [14]. This fungal infection is relatively rare worldwide except in India and Pakistan but highly fatal if inadequately treated. COVID-19-associated mucormycosis (CAM) has been reported from at least 45 countries [15]. India had more than 47,000 cases of mucormycosis reported during the second wave of COVID-19 in May to July 2021 [16]. With a broad-spectrum activity and a low incidence of drug resistance [17], AmB is the first line therapy for invasive mucormycosis and has been commonly used (in more than 88% of patients) to treat CAM [16]. AmB kills fungal cells by pore formation after preferentially binding to ergosterol in the fungal cell membrane or by inducing oxidative damage [18, 19].

Interferon-induced transmembrane proteins (IFITM) 1, 2 and 3 comprise a family of interferon-induced antiviral restriction factors which inhibit the infection of many enveloped viruses including influenza A viruses (IAV) [20–24], flaviviruses [25–28], rhabdoviruses and bunyaviruses [29], human metapneumovirus [30], and human immunodeficiency virus [30–32]. IFITM1 is mainly localized at the plasma membrane, while IFITM2 and IFITM3 are present in endosomes and lysosomes [33]. Opposing effects of IFITM proteins in SARS-CoV-2 infection have been reported. While artificial overexpression of IFITM proteins blocks SARS-CoV-2 infection by relocalizing ACE2, endogenous IFITM proteins are cofactors for efficient SARS-CoV-2 infection [34–36]. Other studies [37–39] showed IFITM1, 2, and 3 block SARS-CoV-2 infection. In addition, human IFITM3 inhibits infection at endosomes but enhances virus fusion at the plasma membrane [37].

AmB has been known to enhance infection of IAV by preventing IFITM3-mediated restriction [40], as well as hepatitis E virus [41], SARS-CoV and SARS-like coronaviruses [42]. Given the models of IFITM3-mediated restriction, it was postulated that the IAV enhancement effect was by increasing membrane fluidity and planarity which would allow viral-host receptor interactions and pore formation. In contrast, AmB is also known to inhibit many viruses such as human coronavirus OC43 [43] human immunodeficiency virus [44], enterovirus 71 [45] and Japanese encephalitis virus [46].

Computer-based methods including *in silico* drug discovery and *in silico* drug screening have been increasingly applied to, and revolutionized many aspects of, the pharmaceutical industry [47]. They enable accelerated drug development as well as drug repurposing. Using *in silico* drug screening, we predicted that AmB and Nystatin had strong interactions with SARS-CoV-2 proteins, indicating these antifungals may strongly modulate or inhibit the virus. In addition, AmB has been proposed as a promising treatment of COVID-19 due to its antiviral activities against many viruses [48]. In support of that, a recent pre-print using Vero E6 and Vero 76 cells reported a clear inhibitory effect of AmB and Nystatin against SARS-CoV-2 [49]. However, as AmB has been shown to have both pro- and antiviral effects, in this study, we aimed to understand the impact of representatives from a range of antifungal classes: Polyene (AmB, Nystatin, Natamycin), Azole (Fluconazole) and Echinocandin (Caspofungin) on SARS-CoV-2 as well as their mechanisms of action. Findings from this study will provide important treatment guidelines for CAM in active COVID-19 cases and may facilitate the development of therapeutics against SARS-CoV-2.

## Results

### Virtual screening to identify AmB and Nystatin as potential drugs against SARS-CoV-2

In late 2020 we utilized an FDA-approved in-house library of 12,000 small molecular drugs, curated by the Rahman group at King’s College London, for virtual screening, to identify the most effective modulators or inhibitors against SARS-CoV-2 proteins (spike, protease 3CLpro, and the nsp15 endoribonuclease). 600 drug candidates were initially selected, based on their free energy of binding, using the docking process with Smina [50, 51]. After refinement, an optimised library of 200 drugs were identified and subjected to Genetic Optimization for Ligand Docking (GOLD) [52, 53]. The best-docked poses for the ligands based on fitness function scores and ligand binding positions were selected and further analysed using a 2D ligand-protein interaction map. Interestingly, both Nystatin and AmB were ranked within the top 10 ligands that interacted strongly with both the nsp15 endoribonuclease and the main protease 3CLPro, making them strong candidates to study their effects on SARS-CoV-2.

### SARS-CoV-2 infection enhanced by AmB and Nystatin

We first tested the effect of AmB, Nystatin, Natamycin, Fluconazole and Caspofungin on SARS-CoV-2 infection by incubating Vero E6 and Calu-3 cells with SARS-CoV-2 at a multiplicity of infection (MOI) of 0.001 and 2-fold serial concentrations of these antifungals. Carboxymethyl cellulose (CMC) supplemented with antifungals was added to the corresponding wells at 2 hours post infection (hpi). Antibody staining of the cells at 24 hpi showed increases in the number of foci / well in the presence of AmB or Nystatin, indicative of viral infection enhancement. The other antifungals tested, showed no signs of infection enhancement. At the optimal concentrations in Vero E6 3.12 µM of AmB and 50 µM of Nystatin, the number of foci were 18- and 16-fold higher than the control wells (Figure 1A). The optimal antifungal concentrations in Calu-3 cells were 2-to 4-fold lower than those in Vero E6 (Figure 1B). The SARS-CoV-2 infection enhancement of AmB and Nystatin was also clearly observed in Huh7 cells with the optimal concentrations similar to those in Vero E6 (Figure 1C). Significant enhancement by AmB and Nystatin was evident with infection by the SARS-CoV-2 XBB 1.5 Omicron variant (Supplementary Figure S1).

**Figure 1.**
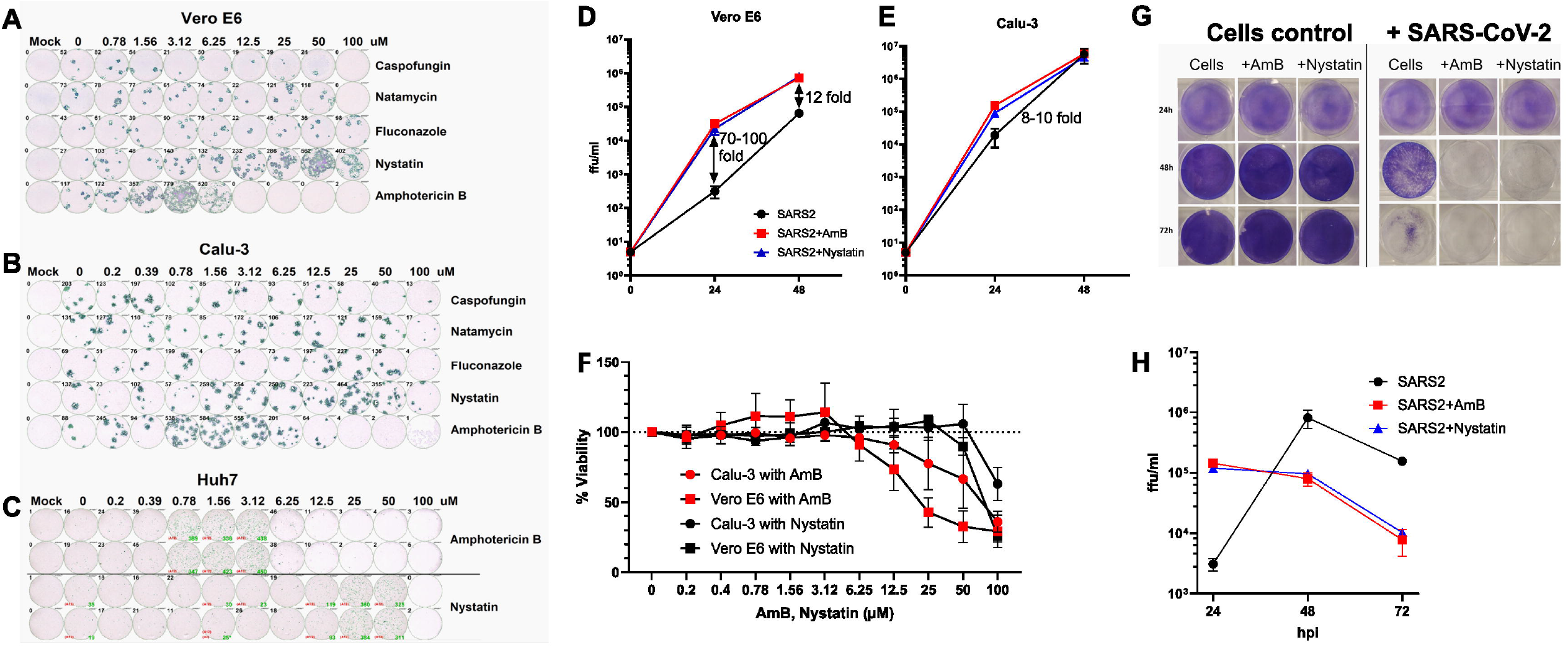
SARS-CoV-2 infection enhancement by AmB and Nystatin. (A-C) Effects of antifungals on SARS-CoV-2 infection in Vero E6, Calu-3 and Huh7 cells. (D-E) Time course of SARS-CoV-2 infection at optimal concentrations of AmB and Nystatin in Vero E6 and Calu-3 cells. (F) MTT assay testing the cytotoxicity of AmB and Nystatin in Vero E6 and Calu-3 cells with the dotted line indicating 100% cell viability. (G) Cell staining with crystal violet shows different levels of cell death (detachment) at 24, 48 and 72 hpi. (H) Titration of supernatant from wells infected with the virus in G.

To understand the magnitude of the infection enhancement, a time course experiment was set up with low MOIs (0.0001 for Vero E6, and 0.00005 for Calu-3) and without CMC. Titration of the supernatant revealed up to 100-fold and 12-fold increase in the viral titres by AmB and Nystatin at 24 and 48 hpi, respectively, in Vero E6 (Figure 1D). Meanwhile, the enhancement effect of these antifungals was about 1 order of magnitude lower in Calu-3 than in Vero E6 (Figure 1E). To determine if drug treatment resulted in cytotoxic effects, an MTT assay was performed. No decrease in cellular viability could be detected for AmB at optimal concentrations (3.12 and 0.78 µM respectively) in VeroE6 or Calu-3 cells; while a minor (10 %) decrease in viability was observed for 50 µM of Nystatin in Vero E6 cells (Figure 1F).

A recent pre-print reported the antiviral activity against SARS-CoV-2 of AmB and Nystatin [49]. In that pre-print, Vero 76 cells were infected with an MOI of 0.1 and treated with a range of concentrations of the antifungals. The supernatant was titrated 48 hpi to determine the effectiveness of AmB and Nystatin. We performed a similar experiment but with optimal enhancement concentrations of 3.12 µM of AmB and 50 µM of Nystatin and the supernatant at 24, 48 and 72 hpi (Figure 1G-H) was titrated. We found that at 24 hpi, these antifungals increased virus titres by 50 - 59-fold compared to the control wells without antifungals. However, at 48 hpi, virus titres in wells with these antifungals were 8 - 10-fold lower than in wells without antifungals. This was due to cell death in wells with antifungals occurring between the two time points while about 75% of cells in untreated wells remained viable at 48 hpi (Figure 1G). At 72 hpi, as expected the virus titer in the control wells also reduced due to cell death at earlier timepoints. We conclude therefore that these antifungals did not have an antiviral effect against SARS-CoV-2. Further experiments to determine the mechanism of enhancement by AmB were carried out.

### AmB promotes SARS-CoV-2 cell entry without affecting cellular gene expression

To determine the effective time points of AmB on SARS-CoV-2 infection, the antifungal was added to the cells at the optimal concentrations (3.12 µM and 0.78 µM to Vero E6 and Calu-3 cells, respectively) at different time points before and after a low MOI infection (MOI 0.01). AmB was removed 1 hour after addition and replaced with fresh medium without AmB. Supernatant harvested at 24 hpi was titrated to assess the extent of viral infection enhancement. Pre-treatment of cells with AmB before virus addition significantly increased SARS-CoV-2 infection in Vero E6 (Figure 2A; 10-fold; p < 0.0001) but not in Calu-3 cells (Figure 2B). Significant enhancement of SARS-CoV-2 infectivity was also observed when AmB was added up to 1 hour after infection to Calu-3 (2.5-fold; p = 0.003), and Vero E6 (12.5-fold; p < 0.0001). In contrast, the addition of AmB after these time points had no considerable effect on SARS-CoV-2 output titres. These findings indicate that the effect of AmB on SARS-CoV-2 infection was due to changes induced by AmB in the host cells rather than to the virus (as shown in the pre-treated Vero E6 cells), and these changes affect an early stage of the virus life cycle.

**Figure 2.**
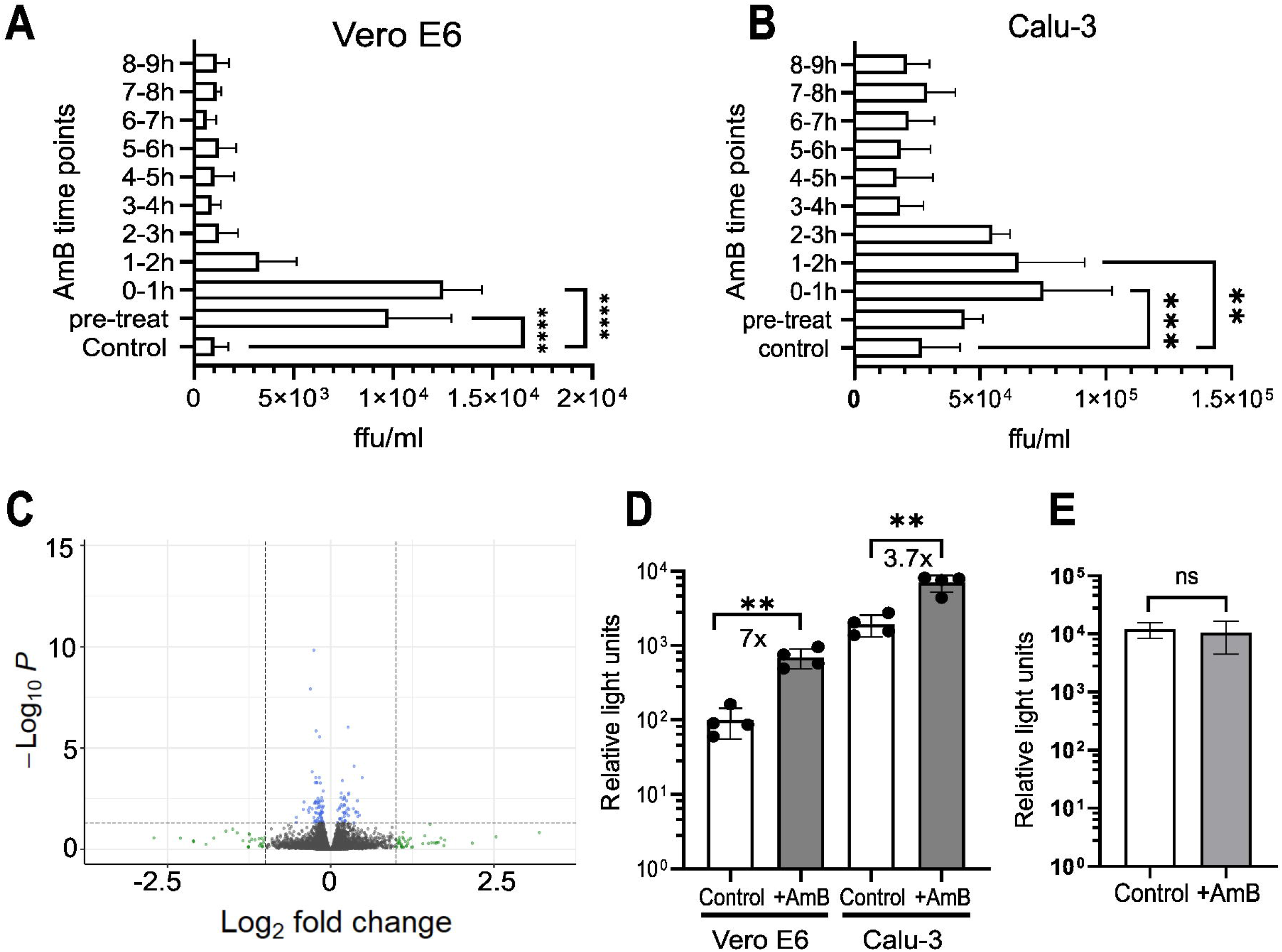
AmB promotes SARS-CoV-2 cell entry without affecting cellular gene expression. (A-B) Vero E6 or Calu-3 cells were treated with AmB at 3.12 µM and 0.78 µM respectively at different time points before (pre-treated) or after infection. AmB was removed 1 hour after addition and replaced with medium without AmB. At 24 hpi, supernatant was harvested and SARS-CoV-2 was titrated. (C) Gene expression changes in Calu-3 cells treated with AmB at 0.78 µM compared to the cells control. The horizontal dashed line represents a q-value of 0.05. The vertical dashed line represents a fold change of 2. (D) Infection of Vero E6 and Calu-3 cells with SARS-CoV-2 pseudovirus in the presence of AmB at 3.2 µM and 0.78 µM, respectively. (E) Vero E6 cells were transfected with pCSFLW plasmid carrying the reporter gene and incubated with or without AmB 3.2 µM. Luciferase assay was performed at 48 hours post transfection (** P < 0.01, *** P < 0.001, **** P < 0.0001, ns non-significant).

We therefore employed RNA-Seq to investigate whether AmB enhanced SARS-CoV-2 cell entry through up- or down-regulation of host genes. However, no significant differences in the gene expression were observed in Calu-3 cells treated for 10 hours with AmB 0.78 µM compared to the untreated control (Figure 2C).

We next used SARS-CoV-2 pseudovirus (replication-deficient lentivirus pseudotyped with the wild-type SARS-CoV-2 spike) to investigate whether AmB promotes viral cell entry. The pseudovirus assay separates viral entry from other steps of the viral infection cycle such as replication. Vero E6 and Calu-3 cells infected with SARS-CoV-2 pseudovirus in the presence of 3.2 µM and 0.78 µM AmB respectively, resulted in substantial increases (7-fold in Vero E6, 3.7-fold in Calu-3 cells) in the luminescence signal compared to the untreated control (Figure 2D). To determine whether AmB enhances the reporter gene (luciferase) expression, we transfected HEK 293T cells with pCSFLW (a plasmid carrying the reporter gene used for SARS-CoV-2 pseudovirus production), and then treated the cells with AmB. The control group did not receive AmB treatment. At 48 hours post transfection, a luciferase assay was performed, showing no statistically significant difference in signals between the treated and untreated wells (Figure 2E). These findings indicated that AmB enhances SARS-CoV-2 pseudovirus cell entry.

### AmB promotes SARS-CoV-2 infection by potentially interfering with the antiviral activity of the IFITM3 protein

We observed that SARS-CoV-2 infection enhancement by AmB was considerably more evident in Vero E6 (for which SARS-CoV-2 mainly enters via endocytosis) than in Calu-3 (for which membrane fusion is the main entry pathway of SARS-CoV-2). Moreover, IAV also enters cells via endocytosis and it was proposed that AmB improves IAV infection by preventing the antiviral effect of IFITM3. A recent study [54] suggested that IFITM3 on the endosome membrane blocks SARS-CoV-2 infection. We therefore tested whether this was also the mechanism through which AmB enhanced SARS-CoV-2 infection. Calu-3 and Huh7 cells were transfected with IFITM siRNAs 24 h before infection with an MOI of 0.001. Treatment with siRNAs reduced the levels of targeted IFITM mRNAs by 65-75% (Figure 3A). Titration of supernatant collected at 48 hpi showed reduced viral titres in wells treated with IFITM1 and IFITM2 siRNAs compared to the siRNA negative control in Calu-3 but not Huh7 cells. In contrast, IFITM3 siRNA had no significant impact on virus titre in Calu-3 cells but remarkably improved the virus titre in Huh7 cells compared to the control (Figure 3B). This, in agreement with previous studies [34, 35], suggested that while IFITM1 and IFITM2 are required for efficient SARS-CoV-2 infection in Calu-3 cells, IFITM3 on the endosome membranes of Huh7 cells inhibits the viral genome release from endosomes in those cells in which SARS-CoV-2 relies mainly on endocytosis for cell entry. To investigate the relationship between IFITM3 siRNA and AmB in Huh7 cells, we treated the cells with IFITM3 siRNA and AmB either individually or in combination before virus infection. Virus titration at 48 hpi revealed a redundant effect of IFITM3 knockdown and AmB (Figure 3C), indicating that both act on the same target. For that reason, AmB plausibly promotes SARS-CoV-2 infection by preventing the antiviral activity of the IFITM3 protein in Huh7 cells. However, it is likely than an alternative mechanism is responsible for the enhanced infection in Calu-3 cells.

**Figure 3.**
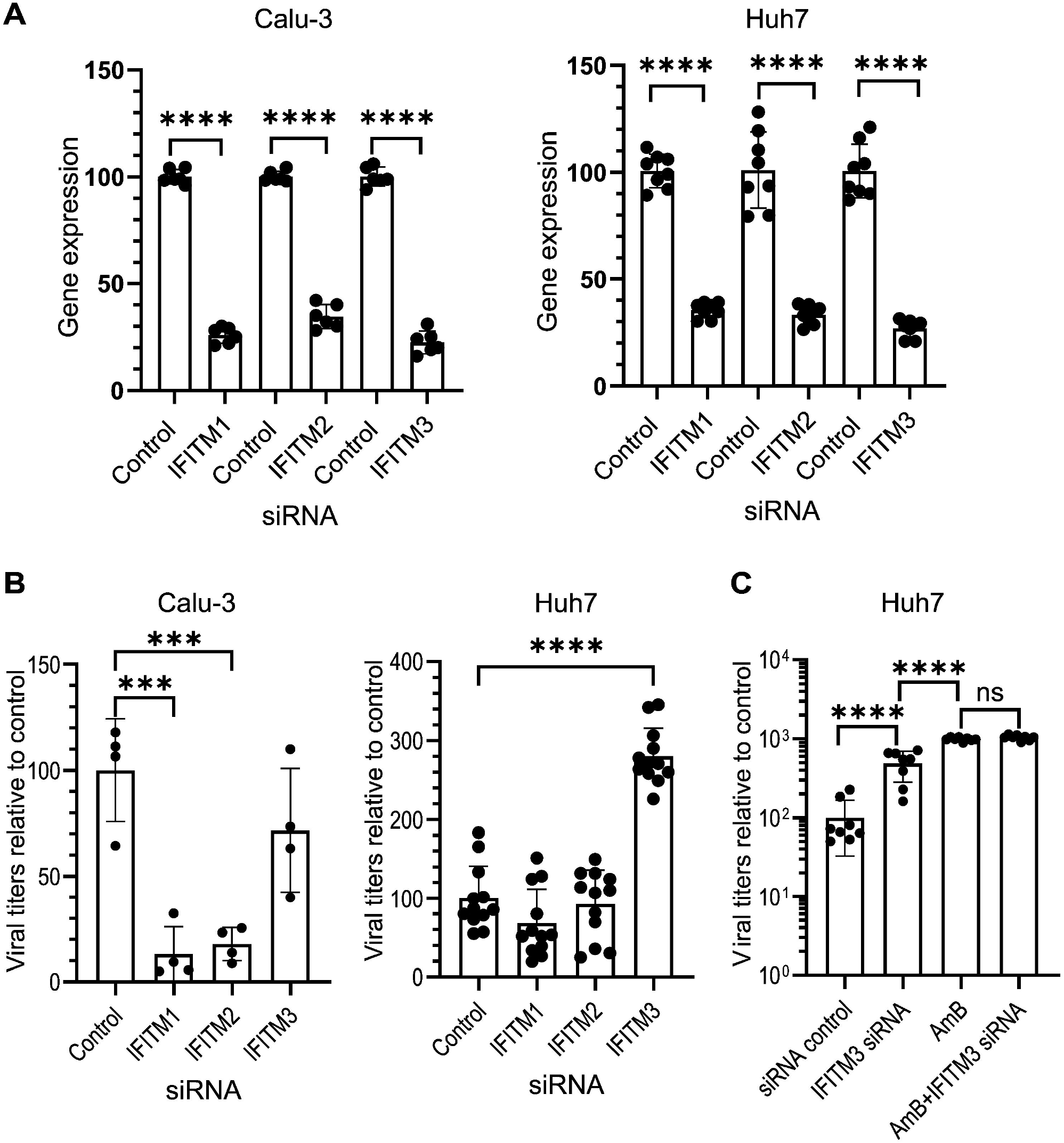
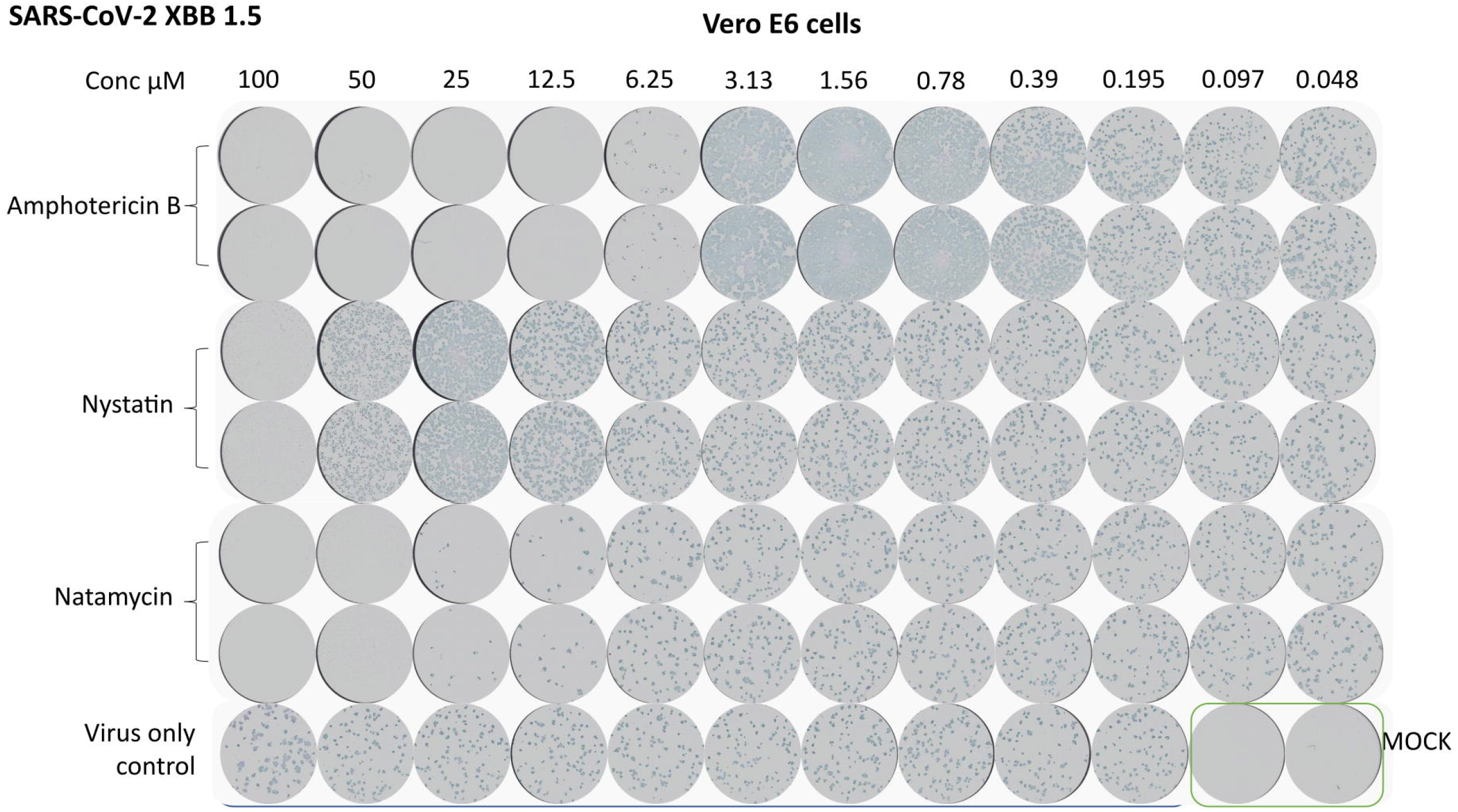
Effects of IFITM siRNAs on SARS-CoV-2 infection in different cell lines. (A) Quantification of IFITM gene expression in Calu-3 and Huh7 cells transfected with siRNA control or IFITM siRNAs. Values were normalized to GAPDH and calculated relative to the control (set to 100%). (B) Extracellular virus titration from Calu-3 and Huh7 cells transfected with siRNA control or IFITM siRNAs before infection with SARS-CoV-2. Values were calculated relative to the control (set to 100%). (C) Extracellular virus titration from Huh7 cells transfected with siRNA control or IFITM3 siRNA before infection with SARS-CoV-2, in the presence or absence of AmB. Values were calculated relative to the control (set to 100%). (*** P < 0.001, **** P < 0.0001).

## Discussion

In this study, using *in silico* screening, we initially predicted that AmB and Nystatin would modulate or inhibit SARS-CoV-2 replication by targeting the viral endoribonuclease and protease. However, our *in vitro* data demonstrated that AmB and Nystatin significantly promote SARS-CoV-2 infection. The level of SARS-CoV-2 enhancement was up to 100-fold higher in Vero E6 cells for which the main viral entry pathway is endocytosis. For Calu-3 cells which primarily support direct membrane fusion for viral entry, up to 10-fold enhancement of infection was observed. Our work furthers previous studies on SARS-CoV-2 [54, 55] and SARS-like coronavirus infection enhancement by AmB [42]. Among these, Peacock and colleagues [54] reported significant SARS-CoV-2 wildtype pseudovirus infection improvement (9-fold) by AmB in 293T-ACE2 cells (low expression level of TMPRSS2) but not in Caco-2 and Calu-3 cells (high TMPRSS2) pre-treated with AmB.

We systematically investigated the mechanism of SARS-CoV-2 infection enhancement by AmB, which has been shown to promote IAV infection by preventing IFITM3-mediated restriction [40]. Using RNAseq and a pseudovirus assay we found that AmB affected the virus entry into cells despite no notable changes in cellular gene expression. Previous studies have used Calu-3 and Caco-2 cells which SARS-CoV-2 enters via direct membrane fusion to avoid the antiviral restriction activity of IFITM3 [54]. In contrast, in Vero E6 and Huh7 cells, SARS-CoV-2 enters cells via endocytosis and IFITM3 on the endosomal membrane could prevent the release of the virus genome into the cytoplasm. IFITM3 knockdown in Huh7 cells enhances SARS-CoV-2 infection. However, in the presence of AmB at its optimal concentration, the effect of IFITM3 knockdown becomes redundant, suggesting that AmB and IFITM3 siRNA target the same pathway or mechanism. The enhancement effect of AmB on SARS-CoV-2 entry into Vero E6 and Huh7 cells, as such, could be attributed to AmB alleviating the effect of IFITM3-mediated restriction, as shown with IAV. This has an implication for the current SARS-CoV-2 dominant variant Omicron, which has evolved to use TMPRSS2 less efficiently and rely more on endocytosis for cell entry. Using AmB in patients infected with Omicron may result in a more significant enhancement of virus infection in tissues in which Omicron enters cells via endocytosis.

A combination of the optimal drug regimen, dose, route of administration and therapy duration, is important not only for treatment success but also for overcoming drug resistance [56]. AmB, the main polyene used as a highly effective antifungal, has a broad-spectrum activity and a low incidence of drug resistance [17]. As such, it has been commonly used to treat CAM [16]. AmB concentrations in pulmonary epithelial lining fluid in patients receiving treatment with liposomal AmB (1.1 – 2.4 µM [57] and 2.3 – 5.3 µM [58]) were reported to be similar to concentrations that resulted in optimal SARS-CoV-2 infection in this study (0.78 - 3.12 µM). Since AmB and Nystatin significantly enhanced SARS-CoV-2 infection, alternative antifungals should be considered when treating CAM in active COVID-19 cases.

Recent studies [59–62] on limited numbers of mechanically ventilated COVID-19 patients showed that these patients may benefit from nebulized AmB as an antifungal prophylaxis to prevent COVID-19 associated pulmonary aspergillosis (CAPA). However, the impact on COVID-19 outcomes was not the main focus of these studies and therefore not reported. Of note, one of these studies [60] revealed active COVID-19 patients with nebulized AmB as antifungal prophylaxis required a significant longer stay in the intensive care units compared to those without antifungal prophylaxis. Randomised trials with larger numbers of participants and monitoring of SARS-CoV-2 in those patients are warranted to confirm the effectiveness of AmB on CAPA prevention as well as its side effects on SARS-CoV-2 infection.

When treating coinfections by different types of pathogens (e.g., viruses and bacteria and/or fungi), off-target effects of drugs on the other pathogen (e.g., effects of anti-mycotics on viruses and vice versa) and on the overall host immune response should be considered. In addition to the virus infection enhancement effect of AmB and Nystatin, other drugs have been shown to have pro-viral activity as an off-target effect. For example, sodium valproate (an anticonvulsant drug) stimulates HIV type 1 [63], human cytomegalovirus [64], measles virus, poliovirus type 1 and coxsackievirus B3 replication [65]. Meanwhile, Chloroquine (a classical antimalarial drug) enhances replication of IAV A/WSN/33 (H1N1) [66], and Cyclosporin A (an immunosuppressant medication) promotes hepatitis E virus infection [67].

The use of computer-based methods for drug discovery and repurposing has shown great promise with the potential to accelerate the drug development process, identify novel drug candidates, and optimize treatment strategies. However, *in silico* screening predictions are not always perfect due to challenges and limitations associated with its application, i.e., lack of sufficient training data, complexity of biological systems, limited understanding of biological mechanisms, unpredictable side effects. This study highlights the crucial validation step when using *in silico* screening for drug repurposing to ensure that the predictions made by these models align with real-world outcomes.

The limitation of this study was that the effects of AmB and Nystatin on SARS-CoV-2 were tested using cultured cell lines, which are likely less representative of the human tissue environment compared to organoids or animal models. However, these findings provide an important insight into the possible off-target effects of antifungals on virus infection which should be investigated further. It also highlights the potential negative impact of commonly used medicines when treating acute viral diseases.

## Materials and methods

### Cells

Vero E6 cells were maintained in Dulbecco’s Modified Eagle Medium (DMEM)-Glutamax supplemented with 10% heat-inactivated foetal calf serum (FBS) and 100 U/ml penicillin, and 100 μg/ml streptomycin. Calu-3 cells were maintained in DMEM-Glutamax mixed with Ham’s F-12 Nutrient Mix (Life Technologies) and supplemented with non-essential amino acids, 10% FBS and 100 U/ml penicillin, and 100 μg/ml streptomycin. For infection steps, DMEM supplemented with 1% FBS was used. Huh7 cells were maintained in DMEM-Glutamax supplemented with 10% FBS, nonessential amino acids, 20 mM HEPES, 100 U/ml penicillin, and 100 μg/ml streptomycin.

### SARS-CoV-2 isolate

Prototype isolate, Victoria/01/202037, received at passage 3 from Public Health England (PHE) Porton Down, passaged in VeroE6/TMPRSS2 cells (NIBSC reference 100978), was used at passage 6. The virus genome sequence was confirmed identical to GenBank MT007544.1, B hCoV-19_Australia_VIC01_2020_ EPI_ ISL_ 406844_ 2020-01-25 by deep sequencing at the Centre for Human Genetics, University of Oxford.

### Virtual screening to identify Nystatin and AmB as potential drugs against SARS-CoV-2

We utilized an FDA-approved in-house library of 12,000 small molecular drugs, curated by the Rahman group at King’s College London, for virtual screening. This process aimed to identify the most effective modulators or inhibitors against three specific targets of the SARS-CoV-2 virus: the interface of the spike glycoprotein (S protein) and the host enzyme ACE2 (PDB ID 6LZG), the main protease 3CLpro (PDB ID 6Y2E), and the nsp15 endoribonuclease (PDB ID 6VWW). It was hypothesised that compounds that strongly interacted within the binding pocket of these ligands would either modulate or inhibit their function. For the virtual screening, we employed AutoDock Smina [50, 51], a tool based on the AutoDock Vina scoring function. Smina was used for blind molecular docking to locate the optimal binding sites on the target proteins, exploring all potential binding cavities.

The docking process with Smina involved default settings, sampling nine ligand conformations through the Vina docking routine’s stochastic sampling method. This approach was used for a large-scale virtual screening of the 12,000-drug library. From this screening, we initially selected drug candidates based on their free energy of binding. This selection was further refined by excluding compounds such as anticancer drugs, low dose but highly potent therapeutics, antibiotics, and drugs from unsuitable therapeutic classes like cardiac glycosides. Subsequently, we applied GOLD [52, 53], to dock these refined drugs to the Smina-identified optimal binding sites of the SARS-CoV-2 targets. GOLD molecular docking was performed to generate a variety of binding orientations and affinities for each mode. The most favourable binding mode for the system was determined by considering the lowest binding free energy. We then selected the best-docked poses for the ligands based on fitness function scores and ligand binding positions. A 2D ligand-protein interaction map was generated using BIOVIA Discovery Studio Visualizer 2021 to further analyse these interactions.

### Cytotoxicity assay

The cytotoxicity of antifungals was tested using the MTT Cell Proliferation Assay Kit (Abcam; catalogue number Ab211091) according to the manufacturer’s instructions. Briefly, 20,000 Vero E6 cells or 70,000 Calu-3 cells were seeded per well in 96-well plates and incubated overnight. Antifungals diluted in a 2-fold serial concentration of 0.1 – 100 µM were added to cells and incubated for 24 hours. Medium was then removed and MTT substrate was added to wells and incubated for 3 hours. Pre-warmed MTT solvent was added to wells to dissolve formazan before measurement of absorbance at OD 590 nm.

### Viral infection enhancement testing

Vero E6 and Calu-3 cells were seeded at 20,000 and 70,000 cells / well, respectively, in 96-well plates, a day before infection. Cells were infected with 50 ffu / well of SARS-CoV-2. Antifungals were diluted in medium and added to wells to obtain the indicated final concentrations. Two hours post infection, CMC supplemented with antifungals was added to wells (100 µl / well). At 24 hpi, plates were fixed with 4% paraformaldehyde and stained with the FB9B monoclonal antibody [68] (kindly provided to us by Alain R. Townsend, University of Oxford, UK) at 1□µg□per ml and secondary antibody conjugated with horseradish peroxidase. TrueBlue substrate was added and the number of foci in each well counted using a CTL Immunospot analyser.

To determine the magnitude of the infection enhancement, one million Vero E6 or Calu-3 in 6-well plates were infected with low MOIs of 0.0001 and 0.00001, respectively, in the presence or absence of AmB or Nystatin at optimal enhancement concentrations. Supernatant was harvested at 24, 48 and 72 hpi and titrated.

To verify the contrast between our findings and those shown by Wasan et. al. [49], a similar experiment was performed. Vero E6 were seeded in 6-well plates at 700,000 cells per well and incubated overnight. Cells were then incubated with or without AmB and Nystatin at optimal enhancement concentrations and infected with an MOI of 0.001. At 24, 48 and 72 hpi, supernatant was titrated and cells were stained with crystal violet as an indirect quantification of cell death.

### Effective time points of AmB

Vero E6 and Calu-3 cells were seeded at 20,000 and 70,000 cells / well, respectively, in 96-well plates. AmB was added to pre-treatment wells at the time of seeding. The following day, cells were infected with SARS-CoV-2 at an MOI of 0.001. AmB was added to wells at indicated time points and removed 1 hour after addition. Titration of the supernatants was performed at 24 hpi.

### Effect of AmB on SARS-CoV-2 pseudovirus entry

Vero E6 cells were seeded at 40,000 cells / well in a 96-well plate and infected with SARS-CoV-2 wild-type pseudovirus at an MOI 0.2 in the presence or absence of 3.2 µM AmB final concentration. At 60 hpi, medium was removed and 100 µl Bright-Glo luciferase substrate (Promega, catalogue number E2620) was added per well. Luciferase signals were measured using a GloMax microplate reader.

To evaluate whether AmB enhances the reporter gene expression, we seeded 15,000 Vero E6 cells per well in a 96-well plate and incubated them overnight. The cells were then transfected with 100 ng of pCSFLW per well using the FuGENE HD transfection reagent (Promega, catalog number E2311), either with or without AmB. At 48 hours post transfection, a luciferase assay was performed as above.

### RNA-Seq

Calu-3 cells were seeded at 400,000 cells / well in a 24-well plate and incubated overnight. Cells were then incubated with or without AmB, 0.78 µM final concentration (4 wells per group). After a 10-hour incubation, medium was removed and RNA was extracted from AmB treated cells and control cells using the Direct-zol RNA Miniprep (Zymo Research, catalogue number R2053). mRNA was isolated from the extracted RNA using the NEBNext® Poly(A) mRNA Magnetic Isolation Module (New England BioLabs, catalogue number E7490S) and libraries were prepared using NEBNext® Ultra™ II RNA Library Prep Kit for Illumina (New England BioLabs, catalogue number E7770S). Libraries were sequenced on a NextSeq device using the 500 / 550 high output v2.5 (150 cycles) kit. Raw output data were filtered to remove low quality reads using TrimGalore (https://github.com/FelixKrueger/TrimGalore) before mapping to the GRCh38.p14 human transcriptome using STAR [69]. Mapped reads were quantified using HTSeq 2.0 [70] before differential gene expression was analysed using edgeR [71].

### IFITM3 knockdown

Huh7 and Calu-3 cells were seeded at 100,000 and 200,000 cells / well, respectively, in 24-well plates and incubated overnight. Cells were then transfected with non-targeting siRNA or siRNAs targeting IFITM genes (Thermo Fisher) at 10 nM final concentration using RNAiMAX (Thermo Fisher, catalogue number 13778100) according to the manufacturer’s instructions. After 24 hours, cells were infected with SARS-CoV-2 at an MOI of 0.001. At 48 hpi, supernatants were harvested for virus titration and cells for qRT-PCR analysis using published primers and probes [34].

### Statistical analyses

T test in GraphPad Prism 10.3.0 was used to compare the means of two groups. For comparison of three groups or more, one-way ANOVA was applied.

## Data Availability

The data that support the findings of this study are available from the corresponding author, [DN], upon reasonable request.

## Acknowledgements

This study was supported by funding from the US Food and Drug Administration Medical Countermeasures Initiative (contract 75F40120C00085). We would like to thank Jack Mellors, Luke Jones and Ola Diebold for critical reading of the manuscript.

## References

1. Bao, L., et al., The pathogenicity of SARS-CoV-2 in hACE2 transgenic mice. Nature, 2020. 583(7818): p. 830–833.

2. Bayati, A., et al., SARS-CoV-2 infects cells after viral entry via clathrin-mediated endocytosis. Journal of Biological Chemistry, 2021. 296: p. 100306.

3. Li, X., et al., Spike protein mediated membrane fusion during SARS-CoV-2 infection. Journal of Medical Virology, 2023. 95(1): p. e28212.

4. Davidson, A.M., J. Wysocki, and D. Batlle, Interaction of SARS-CoV-2 and Other Coronavirus With ACE (Angiotensin-Converting Enzyme)-2 as Their Main Receptor. Hypertension, 2020. 76(5): p. 1339–1349.

5. Koch, J., et al., TMPRSS2 expression dictates the entry route used by SARS-CoV-2 to infect host cells. The EMBO Journal, 2021. 40(16): p. e107821.

6. Jackson, C.B., et al., Mechanisms of SARS-CoV-2 entry into cells. Nature Reviews Molecular Cell Biology, 2022. 23(1): p. 3–20.

7. Dittmar, M., et al., Drug repurposing screens reveal cell-type-specific entry pathways and FDA-approved drugs active against SARS-Cov-2. Cell Reports, 2021. 35(1): p. 108959.

8. Mohamed, M.M. and B.F. Sloane, Cysteine cathepsins: multifunctional enzymes in cancer. Nat Rev Cancer, 2006. 6(10): p. 764–75.

9. Shirato, K., et al., Clinical Isolates of Human Coronavirus 229E Bypass the Endosome for Cell Entry. Journal of Virology, 2017. 91(1): p. 10.1128/jvi.01387-16.

10. Shirato, K., M. Kawase, and S. Matsuyama, Wild-type human coronaviruses prefer cellsurface TMPRSS2 to endosomal cathepsins for cell entry. Virology, 2018. 517: p. 9–15.

11. Meng, B., et al., Altered TMPRSS2 usage by SARS-CoV-2 Omicron impacts infectivity and fusogenicity. Nature, 2022. 603(7902): p. 706–714.

12. Wu, Z. and J.M. McGoogan, Characteristics of and Important Lessons From the Coronavirus Disease 2019 (COVID-19) Outbreak in China: Summary of a Report of 720314 Cases From the Chinese Center for Disease Control and Prevention. JAMA, 2020. 323(13): p. 1239–1242.

13. Guan, W.-j., et al., Clinical Characteristics of Coronavirus Disease 2019 in China. New England Journal of Medicine, 2020. 382(18): p. 1708–1720.

14. Hoenigl, M., et al., The emergence of COVID-19 associated mucormycosis: a review of cases from 18 countries. The Lancet Microbe, 2022. 3(7): p. e543–e552.

15. Özbek, L., et al., COVID-19–associated mucormycosis: a systematic review and meta-analysis of 958 cases. Clinical Microbiology and Infection, 2023. 29(6): p. 722–731.

16. Muthu, V., et al., Epidemiology and Pathophysiology of COVID-19-Associated Mucormycosis: India Versus the Rest of the World. Mycopathologia, 2021: p. 1–16.

17. Vincent, B.M., et al., Fitness Trade-offs Restrict the Evolution of Resistance to Amphotericin B. PLOS Biology, 2013. 11(10): p. e1001692.

18. Mesa-Arango, A.C., L. Scorzoni, and O. Zaragoza, It only takes one to do many jobs: Amphotericin B as antifungal and immunomodulatory drug. Frontiers in Microbiology, 2012. 3(286).

19. Kristanc, L., et al., The pore-forming action of polyenes: From model membranes to living organisms. Biochimica et Biophysica Acta (BBA) - Biomembranes, 2019. 1861(2): p. 418–430.

20. Feeley, E.M., et al., IFITM3 inhibits influenza A virus infection by preventing cytosolic entry. PLoS Pathog, 2011. 7(10): p. e1002337.

21. Klein, S., et al., IFITM3 blocks influenza virus entry by sorting lipids and stabilizing hemifusion. Cell Host & Microbe, 2023. 31(4): p. 616-633.e20.

22. Everitt, A.R., et al., IFITM3 restricts the morbidity and mortality associated with influenza. Nature, 2012. 484(7395): p. 519–23.

23. Bailey, C.C., et al., Ifitm3 limits the severity of acute influenza in mice. PLoS Pathog, 2012. 8(9): p. e1002909.

24. Smith, S.E., et al., Chicken interferon-inducible transmembrane protein 3 restricts influenza viruses and lyssaviruses in vitro. J Virol, 2013. 87(23): p. 12957–66.

25. Brass, A.L., et al., The IFITM proteins mediate cellular resistance to influenza A H1N1 virus, West Nile virus, and dengue virus. Cell, 2009. 139(7): p. 1243–54.

26. Chmielewska, A.M., et al., The Role of IFITM Proteins in Tick-Borne Encephalitis Virus Infection. J Virol, 2022. 96(1): p. e0113021.

27. Wilkins, C., et al., IFITM1 is a tight junction protein that inhibits hepatitis C virus entry. Hepatology, 2013. 57(2): p. 461–9.

28. Narayana, S.K., et al., The Interferon-induced Transmembrane Proteins, IFITM1, IFITM2, and IFITM3 Inhibit Hepatitis C Virus Entry. J Biol Chem, 2015. 290(43): p. 25946–59.

29. Mudhasani, R., et al., IFITM-2 and IFITM-3 but not IFITM-1 restrict Rift Valley fever virus. J Virol, 2013. 87(15): p. 8451–64.

30. McMichael, T.M., et al., IFITM3 Restricts Human Metapneumovirus Infection. J Infect Dis, 2018. 218(10): p. 1582–1591.

31. Diamond, M.S. and M. Farzan, The broad-spectrum antiviral functions of IFIT and IFITM proteins. Nature Reviews Immunology, 2013. 13(1): p. 46–57.

32. Huang, I.C., et al., Distinct patterns of IFITM-mediated restriction of filoviruses, SARS coronavirus, and influenza A virus. PLoS Pathog, 2011. 7(1): p. e1001258.

33. Bailey, C.C., et al., IFITM-Family Proteins: The Cell’s First Line of Antiviral Defense. Annu Rev Virol, 2014. 1: p. 261–283.

34. Prelli Bozzo, C., et al., IFITM proteins promote SARS-CoV-2 infection and are targets for virus inhibition in vitro. Nature Communications, 2021. 12(1): p. 4584.

35. Xie, Q., et al., Endogenous IFITMs boost SARS-coronavirus 1 and 2 replication whereas overexpression inhibits infection by relocalizing ACE2. iScience, 2023. 26(4): p. 106395.

36. Nchioua, R., et al., SARS-CoV-2 Variants of Concern Hijack IFITM2 for Efficient Replication in Human Lung Cells. J Virol, 2022. 96(11): p. e0059422.

37. Shi, G., et al., Opposing activities of IFITM proteins in SARS-CoV-2 infection. Embo j, 2021. 40(3): p. e106501.

38. Xu, F., et al., IFITM3 Inhibits SARS-CoV-2 Infection and Is Associated with COVID-19 Susceptibility. Viruses, 2022. 14(11).

39. Kenney, A.D., et al., Interferon-induced transmembrane protein 3 (IFITM3) limits lethality of SARS-CoV-2 in mice. EMBO Rep, 2023. 24(4): p. e56660.

40. Lin, T.-Y., et al., Amphotericin B increases influenza A virus infection by preventing IFITM3-mediated restriction. Cell reports, 2013. 5(4): p. 895–908.

41. Chew, N., et al., Independent Evaluation of Cell Culture Systems for Hepatitis E Virus. Viruses, 2022. 14(6).

42. Zheng, M., et al., Bat SARS-Like WIV1 coronavirus uses the ACE2 of multiple animal species as receptor and evades IFITM3 restriction via TMPRSS2 activation of membrane fusion. Emerg Microbes Infect, 2020. 9(1): p. 1567–1579.

43. Drayman, N., et al., Drug repurposing screen identifies masitinib as a 3CLpro inhibitor that blocks replication of SARS-CoV-2 in vitro. bioRxiv, 2020.

44. Pleskoff, O., M. Seman, and M. Alizon, Amphotericin B derivative blocks human immunodeficiency virus type 1 entry after CD4 binding: effect on virus-cell fusion but not on cell-cell fusion. J Virol, 1995. 69(1): p. 570–4.

45. Xu, F., et al., Amphotericin B Inhibits Enterovirus 71 Replication by Impeding Viral Entry. Scientific Reports, 2016. 6(1): p. 33150.

46. Kim, H., et al., Antiviral effect of amphotericin B on Japanese encephalitis virus replication. Journal of microbiology and biotechnology, 2004. 14(1): p. 121–127.

47. Lin, X., X. Li, and X. Lin, A Review on Applications of Computational Methods in Drug Screening and Design. Molecules, 2020. 25(6).

48. Al-Khikani, F.H.O., Amphotericin B as antiviral drug: Possible efficacy against COVID-19. Ann Thorac Med, 2020. 15(3): p. 118–124.

49. Wasan, K. and C. Galliano, Determining the antiviral activity of two polyene macrolide antibiotics following treatment in Kidney Cells infected with SARS-CoV-2 bioRxiv, 2022.

50. Koes, D.R., M.P. Baumgartner, and C.J. Camacho, Lessons learned in empirical scoring with smina from the CSAR 2011 benchmarking exercise. J Chem Inf Model, 2013. 53(8): p. 1893–904.

51. Trott, O. and A.J. Olson, AutoDock Vina: Improving the speed and accuracy of docking with a new scoring function, efficient optimization, and multithreading. Journal of Computational Chemistry, 2010. 31(2): p. 455–461.

52. Jones, G., P. Willett, and R.C. Glen, Molecular recognition of receptor sites using a genetic algorithm with a description of desolvation. J Mol Biol, 1995. 245(1): p. 43–53.

53. Jones, G., et al., Development and validation of a genetic algorithm for flexible docking. J Mol Biol, 1997. 267(3): p. 727–48.

54. Peacock, T.P., et al., The furin cleavage site in the SARS-CoV-2 spike protein is required for transmission in ferrets. Nature Microbiology, 2021. 6(7): p. 899–909.

55. Dadonaite, B., et al., A pseudovirus system enables deep mutational scanning of the full SARS-CoV-2 spike. Cell, 2023. 186(6): p. 1263-1278.e20.

56. Silva, L.N., et al., Fungal Infections in COVID-19-Positive Patients: A Lack of Optimal Treatment Options. Curr Top Med Chem, 2020. 20(22): p. 1951–1957.

57. Weiler, S., et al., Pulmonary epithelial lining fluid concentrations after use of systemic amphotericin B lipid formulations. Antimicrob Agents Chemother, 2009. 53(11): p. 4934–7.

58. Moriyama, B., et al., Pharmacokinetics of liposomal amphotericin B in pleural fluid. Antimicrobial agents and chemotherapy, 2010. 54(4): p. 1633–1635.

59. Van Ackerbroeck, S., et al., Inhaled liposomal amphotericin-B as a prophylactic treatment for COVID-19-associated pulmonary aspergillosis/aspergillus tracheobronchitis. Critical Care, 2021. 25(1): p. 298.

60. Melchers, M., et al., Nebulized Amphotericin B in Mechanically Ventilated COVID-19 Patients to Prevent Invasive Pulmonary Aspergillosis: A Retrospective Cohort Study. Crit Care Explor, 2022. 4(5): p. e0696.

61. Rombauts, A., et al., Antifungal prophylaxis with nebulized amphotericin-B in solid-organ transplant recipients with severe COVID-19: a retrospective observational study. Frontiers in Cellular and Infection Microbiology, 2023. 13.

62. Soriano, M.C., et al., Inhaled amphotericin B lipid complex for prophylaxis against COVID-19-associated invasive pulmonary aspergillosis. Intensive Care Medicine, 2022. 48(3): p. 360–361.

63. Moog, C., et al., Sodium Valproate, an Anticonvulsant Drug, Stimulates Human Immunodeficiency Virus Type 1 Replication Independently of Glutathione Levels. Journal of General Virology, 1996. 77(9): p. 1993–1999.

64. Kuntz-Simon, G. and G. Obert, Sodium valproate, an anticonvulsant drug, stimulates human cytomegalovirus replication. Journal of General Virology, 1995. 76(6): p. 1409–1415.

65. Kabiri, M., Motamedifar, M., Akbari, S., The effect of sodium valproate on measles virus, poliovirus type 1 and coxsackievirus B3 replication. Irn J Med Sci 2001. 26: p. 55–61.

66. Wu, L., et al., Chloroquine enhances replication of influenza A virus A/WSN/33 (H1N1) in dose-, time-, and MOI-dependent manners in human lung epithelial cells A549. Journal of Medical Virology, 2015. 87(7): p. 1096–1103.

67. Wang, Y., et al., Calcineurin inhibitors stimulate and mycophenolic acid inhibits replication of hepatitis E virus. Gastroenterology, 2014. 146(7): p. 1775–83.

68. Huang, K.-Y.A., et al., Breadth and function of antibody response to acute SARS-CoV-2 infection in humans. PLOS Pathogens, 2021. 17(2): p. e1009352.

69. Dobin, A., et al., STAR: ultrafast universal RNA-seq aligner. Bioinformatics, 2013. 29(1): p. 15–21.

70. Putri, G.H., et al., Analysing high-throughput sequencing data in Python with HTSeq 2.0. Bioinformatics, 2022. 38(10): p. 2943–2945.

71. Robinson, M.D., D.J. McCarthy, and G.K. Smyth, edgeR: a Bioconductor package for differential expression analysis of digital gene expression data. Bioinformatics, 2010. 26(1): p. 139–40.

